# Abstract Representations of Sensorimotor Transformations in Human Premotor Cortex

**DOI:** 10.64898/2026.08.12.744241

**Authors:** Sanghoon Kang, Juliana E. Trach, Samuel D. McDougle

**Affiliations:** Department of Psychology, Yale University, New Haven, CT, USA; Wu Tsai Institute, Yale University, New Haven, CT, USA; Department of Neuroscience, Johns Hopkins School of Medicine, Baltimore, MD, USA; Kennedy Krieger Institute, Baltimore, MD, USA

## Abstract

Flexible motor control requires representations of context-specific mappings that connect goals to actions. If and how sensorimotor cortical areas encode these mappings is poorly understood. Here, we asked if the human sensorimotor cortex, in addition to encoding movements, also encodes latent mappings between visual goals and movements. Participants learned to adapt wrist movements under two distinct visuomotor transformations, controlling a visual cursor that was either rotated or reflected away from their hand movement. Representational similarity analyses on fMRI data dissociated neural activity patterns related to transformation context, movement direction, and target location. We observed distinct representational profiles within sensorimotor cortex: Premotor cortex activity patterns encoded both movement direction and transformation context, whereas primary motor cortex activity patterns encoded movement direction but not transformation context. In dorsal premotor cortex, stronger transformation context encoding was associated with better task performance. Moreover, in cases where the two transformations converged on the same movement solution, which permitted retrieval of a cached target-response association, participants with stronger behavioral signatures of caching showed greater overlap between transformation and movement representations in dorsal premotor cortex. These findings suggest that premotor cortex flexibly represents abstract sensorimotor transformations linking goals to actions, providing a neural interface between action selection strategies and motor commands.

## Introduction

How does the brain transform intentions into actions? At a basic level, the motor system must convert goals into appropriate movement commands, for example, when planning a reach to a cup on your desk^1–4^. Yet many skilled actions require more than mapping familiar goals onto familiar movements. Humans must also learn to adopt new sensorimotor policies: structured transformations relating goals, movements, and sensory feedback that can be applied across contexts^5–7^.

Consider initially learning to operate a new tool, like a computer mouse. In this example, one needs to learn the structured transformation between movements of the mouse on a horizontal desk and movements of the cursor on a vertical computer screen (e.g., moving the mouse forward moves the cursor upward)^8,9^. Once learned, this mapping can be generalized to new targets, workspaces, postures, and task demands and can also shield learned skills from interfering with each other, allowing one to preserve multiple related skills across contexts. Such flexibility suggests that the motor system represents not only movements, but also sensorimotor transformations that specify how goals should translate into actions, abstracted from the lower-level particulars of specific movements. How the brain learns and represents these sensorimotor policies in a context-sensitive manner remains poorly understood.

One influential account of flexible sensorimotor transformations in the brain focuses on ‘internal models’—neural representations that encode relationships between motor commands and their sensory consequences^10–12^. Internal models are thought to support the gradual and implicit calibration of motor commands in response to sensory prediction errors^13^ and are linked to cerebellar circuits^10,14^. However, rapid learning of novel sensorimotor tasks also depends on deliberate, cognitive strategies^15–17^. In visuomotor adaptation, for instance, people compensate for perturbed visual feedback not only through slow implicit recalibration, but also by explicitly re-aiming their movements to rapidly reduce errors^18–20^. These strategic forms of motor learning allow people to transfer learning of novel visuomotor transformations to a range of new contexts, including novel movement kinematics, perturbation magnitudes, and even across effectors^8,17,21–25^. This flexibility suggests that people can form an explicit mental representation of the underlying sensorimotor transformation itself^23^.

People’s ability to quickly learn and implement abstract, generalizable sensorimotor transformations raises important questions about how the motor system is organized. For example, are generalizable sensorimotor transformation rules represented within sensorimotor cortex, where they could directly bridge goals and movements? Several lines of evidence point to the premotor cortex as a potential interface between abstract goals and movements^26–28^. For example, premotor regions contain effector-independent movement information^29–32^, encode upcoming actions and structure in learned motor sequences^33,34^, and exhibit function-based action coding (e.g., show overlapping neural activity patterns for curling the toes and grasping with the fingers)^31,35^. Moreover, in studies of cognitive control that require the maintenance of different, sometimes competing, stimulus-response rules, the premotor cortex has been highlighted as one of several frontal cortex regions that can maintain abstract contextual information to guide action selection^36,37^. These findings point to representational codes in motor areas that are more abstract than low-level motor commands, yet are still closely tied to action planning and execution.

Here we used functional neuroimaging to investigate representations of sensorimotor transformations in the human sensorimotor cortex. Participants learned to guide a visual cursor using hand movements under two distinct visuomotor transformations: rotation and mirror reflection. Our task design allowed us to dissociate neural encoding of specific movements versus encoding of transformation context abstracted across movement directions and target locations. We found that premotor cortex, but not primary motor cortex, reliably carried a multiplexed representation of both motor commands and the abstracted transformation context.

## Results

Human participants made hand movements under two distinct visuomotor transformations — rotation and reflection (**Fig. 1a**). Under visuomotor rotation (Rotation), participants’ visual feedback (a circular cursor) was rotated 40° clockwise relative to the direction of their movements. In contrast, under visuomotor reflection (Reflection), the visual feedback was reflected across the y-axis of the workspace (see *Methods* for further details). Participants were explicitly cued about the current transformation context at the beginning of each block (**Fig. 1b**). Visual feedback was delayed to isolate strategic motor learning and limit implicit motor adaptation^38^. One of four possible target locations was presented each trial, with two of the target locations requiring unique movement solutions across the transformation contexts (‘unmatched’ trials) and the other two requiring the same movement solution (‘matched’ trials; **Fig. 1c**). This design allowed us to examine the neural representation of the transformation context, while controlling for visual information (i.e., target location) and movement kinematics.

**Figure 1.**
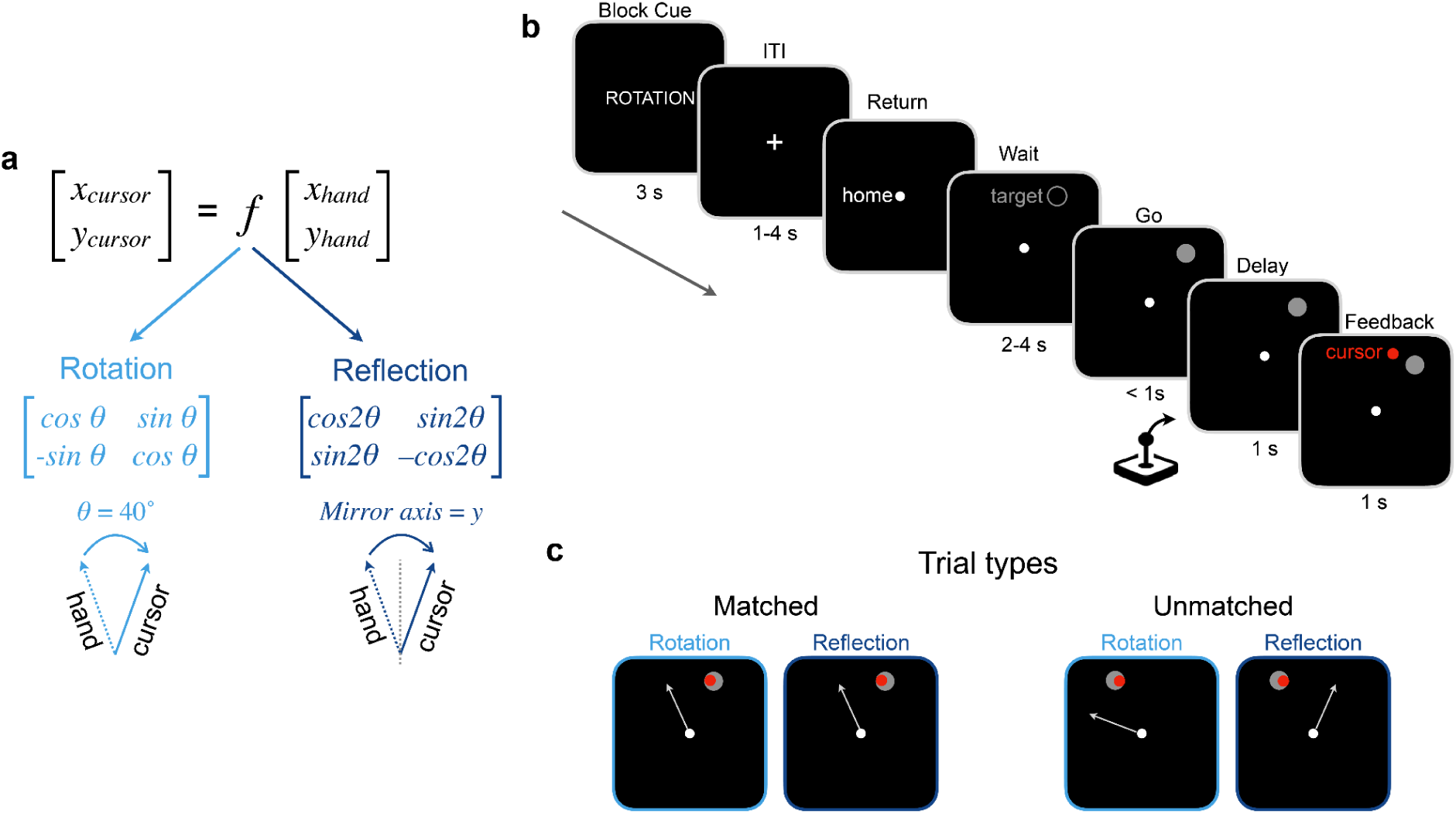
Task design. **a)** Schematic of experiment logic – learning an abstract transformation (e.g., Rotation, Reflection) allows one to map from goals to actions across different transformation contexts (function *f*). Our task implemented two visuomotor transformations: a 40° clockwise rotation of visual feedback and a mirror reflection of feedback across the y-axis. **b)** Trial structure. Blocks began with an explicit instruction cue specifying the currently active transformation. Trials were separated by a jittered ITI, and began with participants putting the joystick in the home position. They then saw a hollow target appear (gray circle) and waited through a jittered delay, before receiving a go cue (target filling) prompting them to move the joystick in a swift flicking motion. Visual feedback of the cursor’s endpoint position was delayed for one second, and then displayed for one second. **c)** Of the four equally probable target locations, two required the same movement in both transformation contexts to successfully contact the target with the cursor (‘matched’ trials; one of the two matched-trial target locations is shown on the left) and two required different movements across transformation contexts (‘unmatched’ trials; one of the two unmatched-trial target locations is shown on the right).

### Behavioral results show comparable learning of rotation and reflection transformations

Participants successfully learned the two transformations and reached a performance asymptote during the tutorial phase (**Fig 2a-c**). Post-training, their average absolute initial angular movement error was modest (Rotation: 17.22°+/-4.58; Reflection: 17.23°+/-3.70). Average error was not significantly different across Rotation versus Reflection conditions and different timepoints (e.g. initial vs. endpoint) of movement (absolute initial error: *t*(26) = -.02, *p* = .98; signed initial error: *t*(26) = -.19, *p* = .85; absolute endpoint error: *t*(26) = .08, *p* = .94; signed endpoint error: *t*(26) = -.95, *p* = .35), showing that both transformations were learned, and that both conditions were of comparable difficulty at the time of scanning.

**Figure 2.**
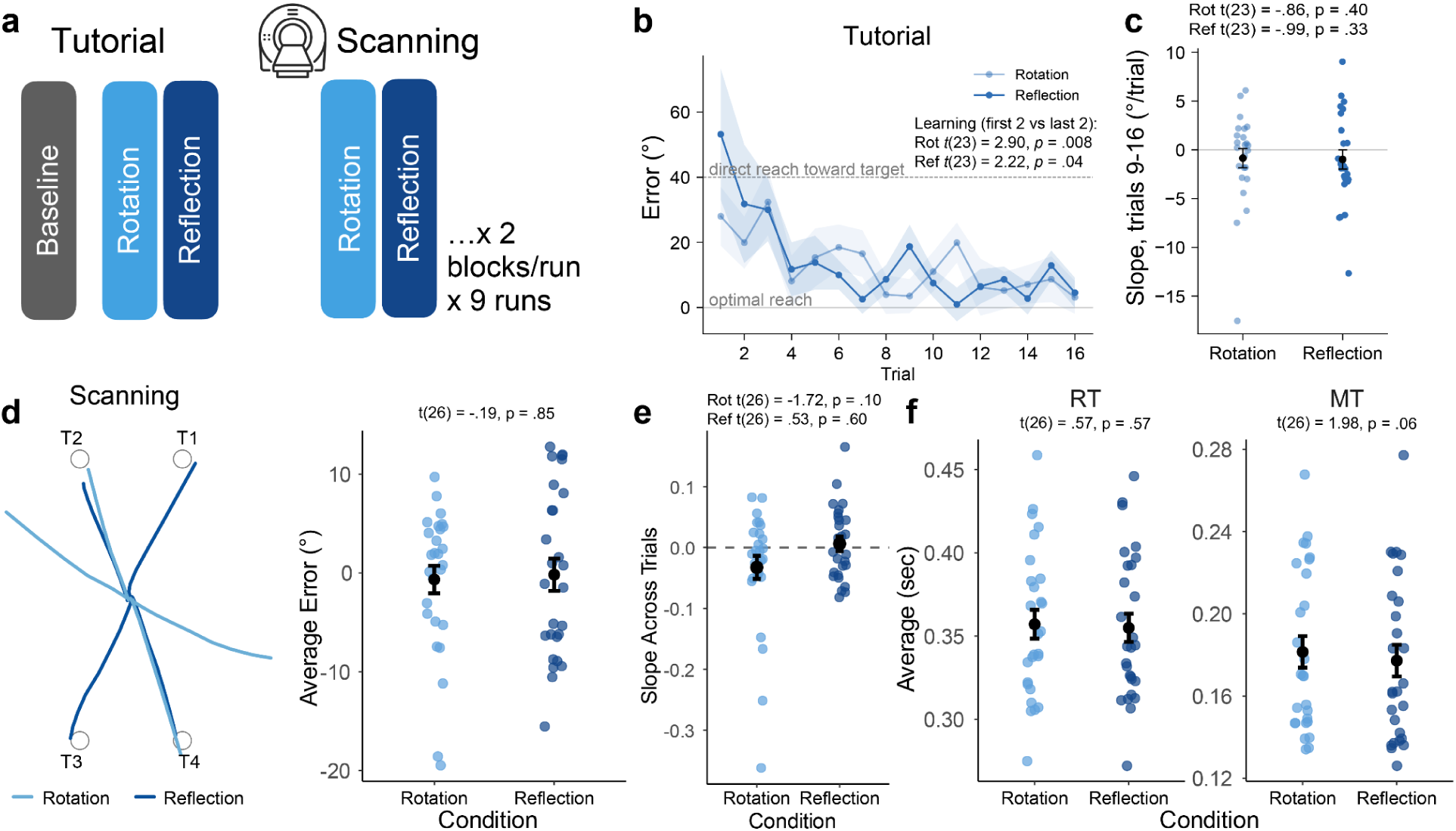
Behavioral results. **a)** Task schedule. Participants were trained (tutorial phase) before undergoing functional scanning. **b)** Average initial heading error for the Rotation and Reflection conditions during the tutorial learning phase. Errors in this figure were standardized to conform to a common direction of perturbation compensation. The inset shows statistical quantifications of learning from the first to the last two trials of each block. **c)** Learning slopes computed on error time courses for the second half of each tutorial block. Inset shows statistical tests of learning slopes. We note that three subjects are omitted from these plots due to an error in saving their tutorial data. **d)** Left: An example participant’s reaching trajectories during scanning across the eight distinct trial types (2 transformations X 4 target locations). Right: Average signed movement error during scanning across participants. **e)** Learning slopes across the scanning runs were not significantly different from zero, consistent with asymptotic performance (see *Results*). **f)** Reaction times (RTs) and movement times (MTs) were not significantly different across transformation contexts. Error shading and error bars represent 1 S.E.M.

To further confirm that participants were successfully trained during the tutorial, we performed a regression analysis on their errors across the scanning session to derive a learning slope (**Fig. 2e**). No significant slope emerged in either transformation condition (Reflection: *t*(26) = .53, *p* = .60; Rotation: *t*(26) = -1.72, *p* = .10), confirming that participants’ accuracy was stable during scanning. Participants also showed comparable reaction times (RTs) and movement times (MTs) across the conditions (**Fig. 2f**; RT: *t*(26) = .57, *p* = .57; MT: *t*(26) = 1.98, *p* = .06).

### Neural pattern analyses reveal distinct representational profiles in premotor versus primary motor areas

We adopted a representational similarity analysis (RSA) multiple regression approach (**Fig. 3a**) to model neural activity patterns as a function of different *a priori* task variables in different brain regions. To do this, we derived a cross-validated empirical representational similarity matrix (RSM) by estimating the Spearman correlation between neural activity patterns across trial types. Then, we defined four predictors in this analysis corresponding to the transformation context (Rotation versus Reflection), the movement direction required to contact the target with the cursor, the visual target location, and the unique trial type (eight in total). By modeling the cross-validated empirical RSM with the four predictors in a multiple regression, we could detect unbiased, predictor-specific information in measured BOLD activity patterns across brain areas (see *Methods*).

**Figure 3.**
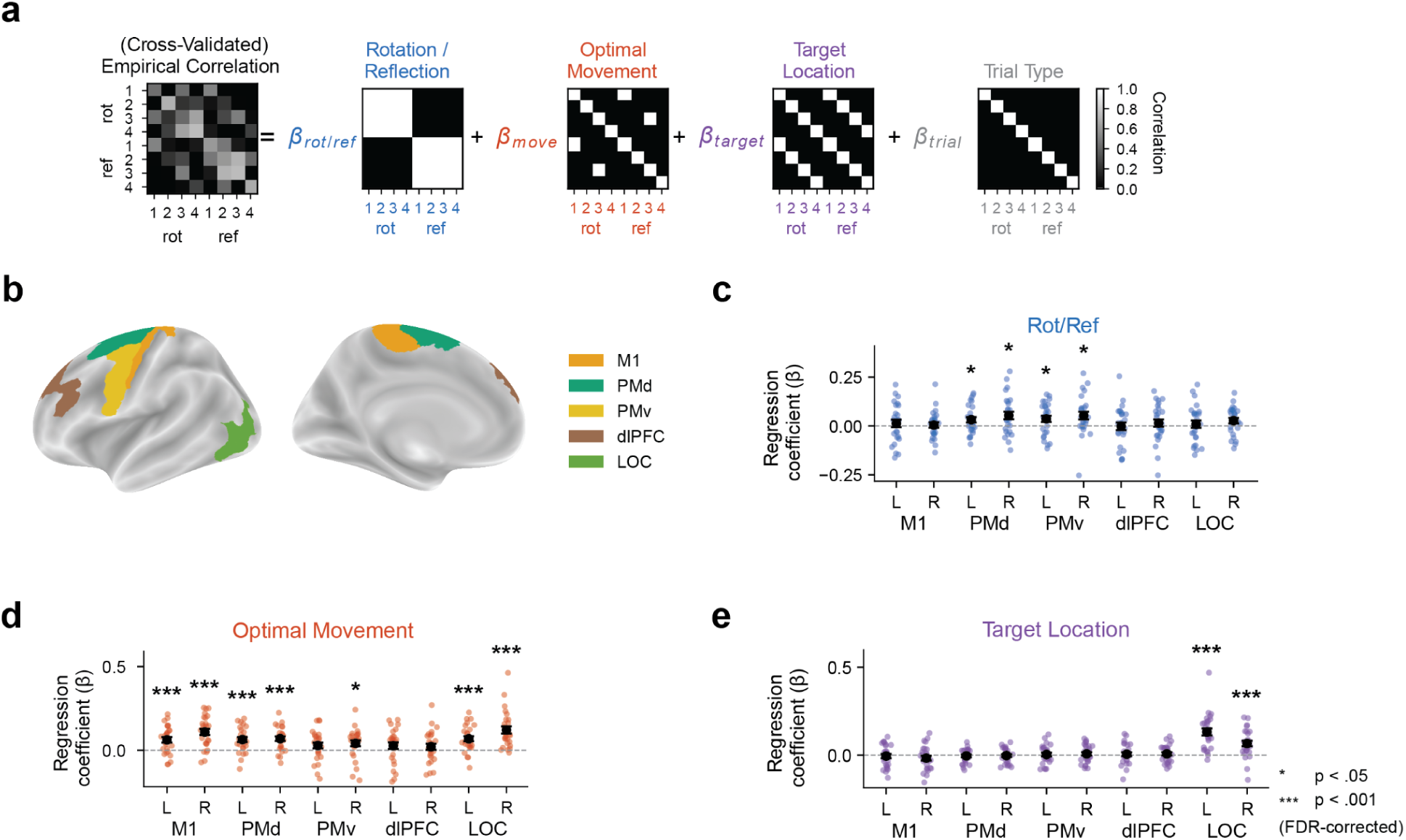
RSA results. **a)** Analysis logic of the RSA multiple regression approach. Hypothesized representational similarity structure for different task variables was regressed against cross-validated empirical neural pattern similarity matrices in multiple anatomical ROIs (see *Methods* for additional details). **b)** *A priori* anatomical ROIs, displayed on the cortical surface. **c-e)** Regression coefficients for **c)** transformation context, **d)** optimal movement, and **e)** visual target location. Asterisks connote significant FDR-corrected t-test results. Error bars = 1 S.E.M. M1 = primary motor cortex; PMd = dorsal premotor cortex; PMv = ventral premotor cortex; dlPFC = dorso-lateral prefrontal cortex; LOC = lateral occipital cortex.

We selected *a priori* anatomical regions of interest (ROIs; **Fig. 3b**) in bilateral premotor cortex (dorsal and ventral; PMd and PMv) and primary motor cortex (M1) to examine how abstract transformation context versus motor commands might be represented across these key sensorimotor cortical regions. We tested additional bilateral prefrontal ROIs in the dorso-lateral prefrontal cortex (dlPFC) as potential neural correlates of putative rule maintenance signatures^39^, and bilateral high-level visual cortex (lateral-occipital cortex, LOC) was selected as a likely neural correlate of target location^40^.

Premotor areas, both dorsal and ventral, showed abstract transformation encoding (**Fig 3c**): Regression weights for the transformation context were significantly positive in bilateral dorsal and ventral premotor cortex (t-tests on regression weights against zero: left dorsal: *t*(26) = 2.28, *p_uncorrected_*= .02, *p_fdr_* = .04; right dorsal: *t*(26) = 2.84, *p_uncorrected_* = .004, *p_fdr_* = .04; left ventral: *t*(26) = 2.26, *p_uncorrected_*= .02, *p_fdr_* = .04; right ventral: *t*(26) = 2.75, *p_uncorrected_* = .005, *p_fdr_* = .04). In contrast, transformation context encoding was not reliable in M1 (left: *t*(26) = .68, *p_uncorrected_* = .25, *p_fdr_* = .33; right: *t*(26) = .34, *p_uncorrected_* = .37, *p_fdr_*= .42). Moreover, we did not see evidence for transformation encoding in our *a priori* prefrontal ROIs (left dlPFC: *t*(26) = -.08, *p_uncorrected_*= .53, *p_fdr_* = .53; right dlPFC: *t*(26) = .78, *p_uncorrected_*= .22, *p_fdr_* = .32), nor in visual cortex (left LOC: *t*(26) = .50, *p_uncorrected_* = .31, *p_fdr_* = .38; right LOC: *t*(26) = 1.98, *p_uncorrected_* = .03, *p_fdr_*= .06). Together, these results suggest that of our *a priori* ROIs, premotor areas uniquely represented sensorimotor transformations abstracted from movement.

We observed reliable encoding of the optimal movement direction across multiple regions (**Fig 3d**). M1 showed significant encoding of movement directions (left: *t*(26) = 3.84, *p_uncorrected_* < .001, *p*_fdr_ < .001; right: *t*(26) = 6.00, *p_uncorrected_*< .001, p_fdr_ < .001), as well as bilateral PMd (left: *t*(26) = 4.46, *p_uncorrected_* < .001, *p_fdr_*< .001; right: *t*(26) = 5.12, *p_uncorrected_* < .001, *p_fdr_* < .001) and the right PMv (*t*(26) = 2.37, *p_uncorrected_* = .01, p_fdr_ = .02; left: *t*(26) = 1.67, *p_uncorrected_* = .05, *p_fdr_* = .06). Bilateral LOC also showed encoding of movement direction (left: *t*(26) = 4.57, *p_uncorrected_* < .001, *p_fdr_* < .001; right: *t*(26) = 5.70, *p_uncorrected_* < .001, *p_fdr_* < .001), which could reflect retinotopic shifts in target location when people direct their gaze toward the planned movement direction^41^. As expected, the visual cortical ROIs showed significant encoding of the target location (**Fig 3e**; left: *t*(26) = 6.60, *p_uncorrected_*< .001, *p_fdr_* < .001; right: *t*(26) = 4.28, *p_uncorrected_*< .001, *p_fdr_* < .001), but no other region showed this effect (*ps_uncorrected_* > .21, *ps_fdr_* > .60).

To statistically test a representational distinction between premotor and M1 ROIs, we performed a difference-of-differences analysis: in each ROI, we computed the difference in coefficient values between transformation context and movement direction, and compared the results between premotor versus M1 regions (ROI x coefficient-type interaction: all *ps* < .05). This result confirmed that transformation context and movement information were weighted differently in premotor versus M1 regions, supporting a multiplexed code of context and movement in premotor regions and a movement-only code in M1. This premotor-versus-M1 distinction was stronger in the right hemisphere when comparing PMd to M1 (ROI x hemisphere interaction: *F*(1, 26) = 7.98, *p* = .009) and marginal when comparing PMv to M1 (*F*(1, 26) = 3.58, *p* = .07).

Finally, we found that encoding of transformation context could not be solely explained via trial-type specific conjunctive representations of context and movement in most ROIs: specific trial type coefficients were not significantly above zero (all *ps_uncorrected_* > .09, *ps_fdr_* > .27 except for left LOC at *t*(26) = 3.69, *p_uncorrected_* < .001, *p_fdr_* = .008). Moreover, the full RSA model that included transformation context consistently outperformed a model that did not (**Fig. S1**). Taken together, our RSA results (**Fig. 3**) suggest that bilateral PMd and right PMv simultaneously encoded sensorimotor transformation context and lower-level movement kinematics, while primary motor cortex only encoded the latter.

We performed additional control analyses to confirm that our results genuinely reflected the neural representation of transformation context. First, we investigated potential univariate differences in premotor areas between transformation contexts. Even though our correlational approach should not be sensitive to univariate differences in premotor activation across the two transformation contexts, univariate differences could still influence our interpretation of the results. However, in a control whole-brain GLM analysis (see *Methods*), we found no significant clusters in or near premotor areas in a univariate contrast between Rotation and Reflection trials (**Fig. S2**). Second, we controlled for lower-level motor differences across transformation contexts: While average motor error was similar across the conditions and not significantly different from zero (**Fig. 2d**), suggesting that subjects generally reached in the optimal movement direction, within-subject differences in movement direction and variability across conditions could still drive neural differentiation of the two transformation contexts in the RSA. That is, our evidence for sensorimotor transformation encoding (**Fig. 3c**) could still be subtly confounded with unmodeled lower-level motor variables. To test this, we re-ran our RSA analyses but using participants’ actual average movement directions in each trial type rather than the optimal movement direction. We still observed significant transformation context encoding in all four premotor ROIs (all *ps* < .05), but not in M1 (all *ps* > .19).

Third, we asked if variability in movement feedback (e.g., observed cursor directions) could be confounding our key premotor results. We performed a behavioral RSA-style analysis^42^ where we first computed initial cursor direction similarity matrices across trial types and then took the same regression approach as in the neural RSA (**Fig. S3**). Then, we used this model’s regression weights for transformation context to model the neural regression weights for the same factor. That is, we examined the relationship between regression weights derived from the neural and behavioral (observed movement error) RSAs to assess whether transformation-context information estimated from the neural RSA can be explained by information derived from the behavioral data. As a result, we found that the behavior-based transformation-context betas were not significantly related to neural transformation-context betas across all ROIs tested. Further, the positive transformation-context effects in premotor areas remained robust even when accounting for these behavioral effects. This analysis supports our conclusion that the results depicted in **Fig. 3c** reflect neural correlates of the abstracted transformation context rather than variation in motor behavior or feedback.

### MDS suggests encoding of transformation context in left PMd is distinct from movement encoding, and transformation encoding covaries with task performance

Our neural RSA results pointed to simultaneous encoding of abstract transformation context and movement direction in premotor areas. How can we think about this finding representationally? To visualize the representational geometry in our regions of interest, we performed multi-dimensional scaling on the group-averaged representational similarity matrices, focusing on the left (contralateral to the performing hand) premotor versus primary motor cortex. As depicted in **Fig. 4a**, left PMd appeared to show distinct coding dimensions roughly corresponding to movement direction and transformation context. Meanwhile, left primary motor cortex representations were clustered around movement directions without showing sensitivity to transformation context (**Fig. 4b**). Additional MDS visualizations across the other anatomical ROIs and hemispheres are depicted in **Fig. S4**.

**Figure 4.**
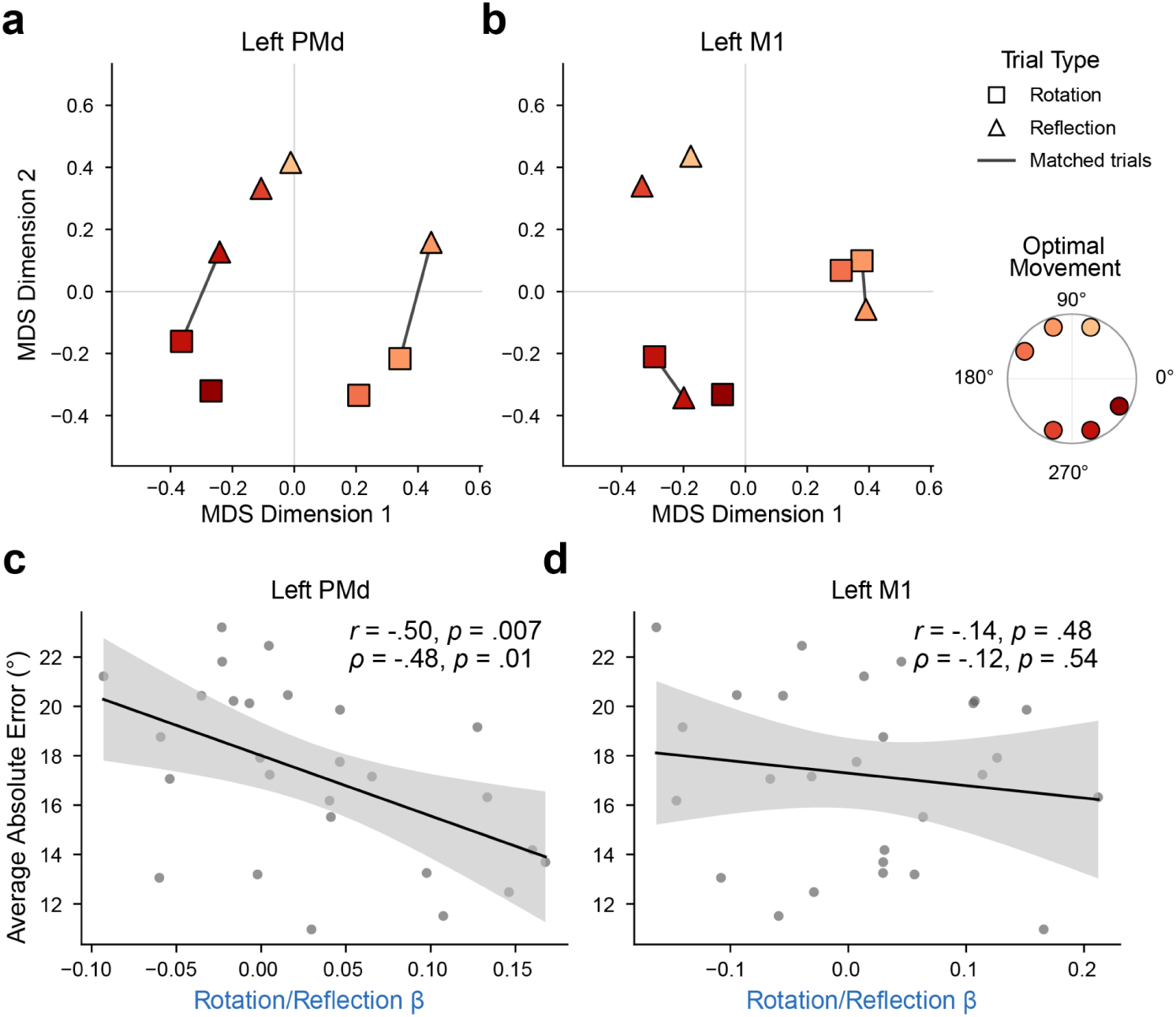
MDS visualizations and brain-behavior correlations. a-b) Results of a multi-dimensional scaling analysis on group-averaged empirical representational dissimilarity matrices, shown for contralateral (left) **a)** PMd and **b)** M1 (see *Methods* for additional analysis details). **c-d)** Brain-behavior correlations (using both *r_pearson_* and *ρ_spearman_*) between abstract transformation context coefficients (see Fig. 3) and overall task performance, for contralateral (left) PMd and M1. Insets: correlation statistics. Error shading = 95% C.I.

We next asked if premotor encoding of sensorimotor transformation context covaried with performance in the task. In a between-subjects brain-behavior correlation analysis, we found that the effect size of abstract transformation encoding in left PMd significantly covaried with overall motor error (**Fig. 4c**; *r_pearson_* = -.50, *p* = .007; *ρ_spearman_*= -.48, *p* = .01), an effect that was not reliable in nearby primary motor cortex (**Fig. 4d**; left: *r_pearson_* = -.14, *p* = .48; right: *r_pearson_* = -.38, *p* = .05). These correlations were also not reliable in the ventral premotor cortex (left: *r_pearson_* = -.12, *p* = .55, right: *r_pearson_* = .13, *p* = .52) and the right dorsal premotor cortex (*r_pearson_* = -.27, *p* = .17). These results suggest that abstract contextual information in PMd is linked to motor performance in our task.

### Evidence for ‘caching’ of stimulus-response policies on trials with matched movement goals between contexts

Our task design included a manipulation (**Fig. 1c**) where one class of trials required the same movement solution across transformation contexts (‘matched trials’) and another class of trials required unique movement solutions across transformation contexts (‘unmatched trials’). This manipulation allowed us to pose another question: when two distinct transformations point to the same movement goal (matched trials), are their neural representations further pulled apart to avoid interference, or might transformation information be reduced due to the implementation of a simpler cognitive strategy (**Fig. 5a**)? This question was inspired by behavioral work showing that visuomotor learning can be supported by distinct cognitive strategies: explicit use of a sensorimotor mapping versus the ‘caching’ of direct stimulus-response associations following sufficient practice^23,43^.

**Figure 5.**
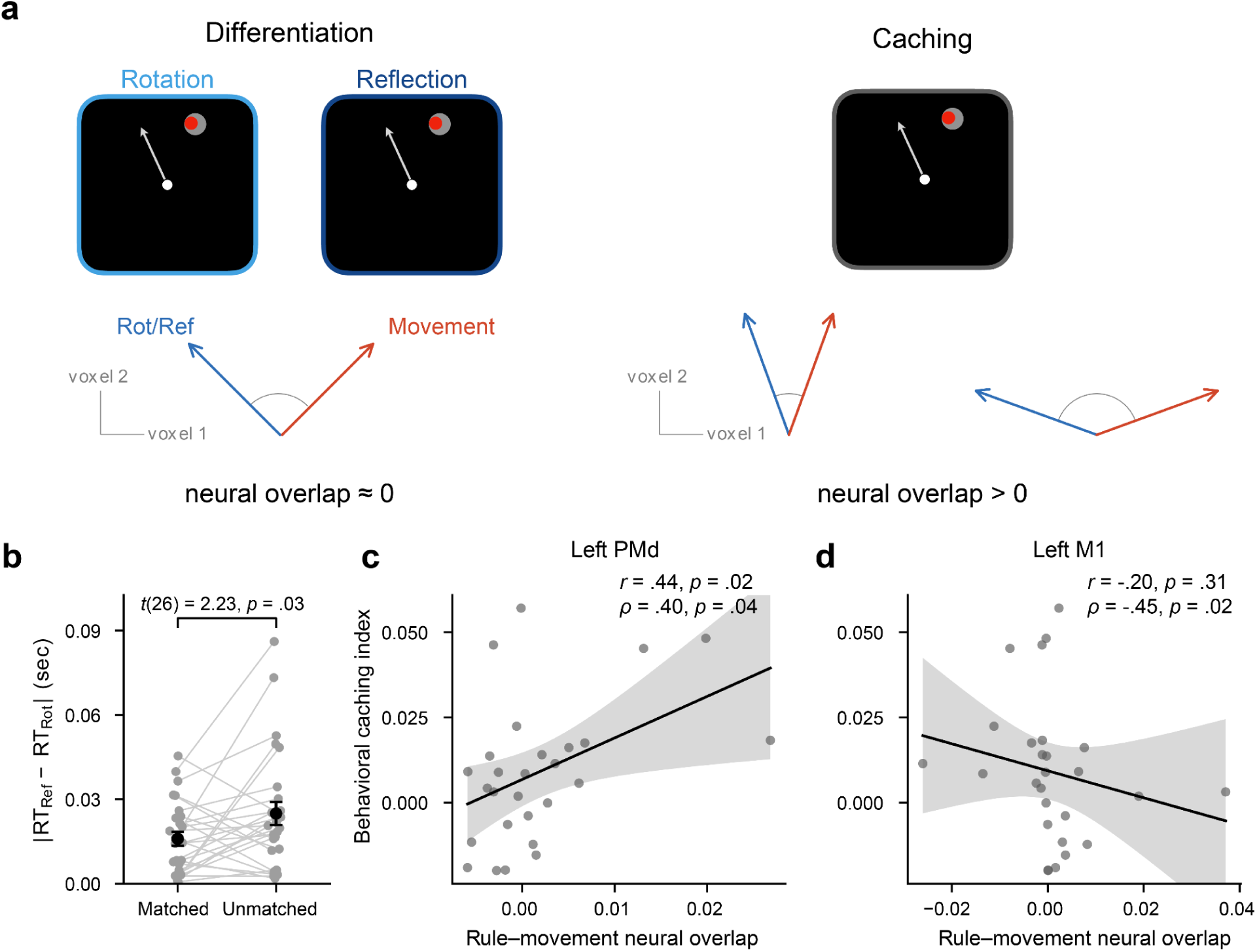
Behavioral and neural evidence for stimulus-response ‘caching’. **a)** Schematic describing two potential strategies on matched trials, where both transformation contexts specified the same movement. A ‘Differentiation’ (left) strategy implies that transformation context is maintained despite the same movement being required, while a ‘Caching’ (right) strategy implies that a stimulus-response association is executed in a context-insensitive manner. Vector schematics represent how these two strategies might relate to neural representational geometry in different ROIs: when the two opposing contexts are distinguished (differentiation), context and movement representations should be orthogonalized, or show no overlap; when both contexts specify the same movement and context can thus be ignored (caching), context and movement representations should show greater overlap (see *Methods*). **b)** Absolute differences in response time (RT) across the two transformation contexts. This metric shows reduced cross-context differences in behavior on matched relative to unmatched trials, consistent with a caching strategy. **c)** Correlations between the behavioral caching effect (the subtraction of matched versus unmatched trials from panel b) and neural overlap of context and movement-related activity pattern components in left PMd. **d)** Same as c, but in left M1. Spearman (rank-based) correlation results are shown in addition to Pearson correlation results to control for apparent outlier values. Error shading represents 95% CI.

We reasoned that matched trials may show evidence for this caching strategy, as they require the same motor response regardless of the current transformation context. Even though overall RTs were comparable between Rotation and Reflection conditions, within-subject RTs were variable such that subjects displayed different reaction times in different conditions (**Fig. 2f)**. We took advantage of this variability to measure caching behaviorally by measuring the extent to which people’s behavior converged across contexts in the matched trials, where the lower-level features of the task (target location and correct reach direction) are equated. Specifically, we examined if the (absolute) difference in RT between contexts was reduced in matched trials relative to unmatched trials. A positive difference in this analysis would suggest that, on matched trials, people implemented a stereotyped stimulus-response strategy across contexts rather than the context-specific rule. Indeed, differences in RT between the Rotation and Reflection conditions were squelched on matched trials relative to unmatched trials (*t*(26) = 2.23, *p* = .03), consistent with caching (**Fig. 5b**). RT was also overall faster on matched trials (*t*(26) = 2.54, *p* = .02), likely because people were provided more opportunities to retrieve putatively cached movements in matched relative to unmatched trials.

Suggestive evidence for caching in the neural data can be seen in **Fig. 4a**, where, in PMd, representational distances across transformation conditions appeared to be reduced in matched versus unmatched trials. However, this is a pooled group-level result. To measure neural evidence of a caching effect and link it to behavioral signatures at the individual level, we computed the ‘overlap’ between isolated context-versus movement-related components of the measured neural patterns in left PMd (**Fig. 5a**; see *Methods*). Increased overlap would reflect one potential neural consequence of behavioral caching on matched trials: caching on matched trials would, in principle, make the contribution of transformation context information be primarily driven by unmatched trials, which entangle transformation context and movement direction. Indeed, this neural correlate of caching was positively correlated with our behavioral measure of caching across subjects in left PMd (**Fig. 5c**; *r_pearson_* = .44, *p* = .02; *ρ_spearman_* = .40, *p* = .04) but not in left M1, where it was non-significant or negative (**Fig. 5d**; *r_pearson_* = -.20, *p* = .31; *ρ_spearman_*= -.45, *p* = .02).

### Transformation context encoding in additional cortical and subcortical regions

Next, we asked which areas beyond our *a priori* anatomical ROIs might also encode information related to sensorimotor transformation context. We approached this exploratory question in two ways. First, we examined neural activity patterns in a second set of candidate anatomical sensorimotor ROIs: the superior parietal lobule (SPL), which has been linked to the encoding of novel sensorimotor mappings^44–46^, and primary somatosensory cortex (S1), known to be involved in the retention of new motor memories^47^. We also examined the hippocampus given its role in context-dependent decision-making^48^. We observed (right-lateralized) transformation context encoding in these three additional ROIs (**Fig. S5**). Unlike contralateral dorsal premotor cortex (**Fig. 4c**), transformation encoding in these areas did not correlate with task performance (all *ps* > .25).

Second, we performed an exploratory whole-brain RSA regression searchlight analysis (see *Methods*) to visualize broader network-level effects (**Fig. 6; Fig. S6**). Searchlight results along the sensorimotor hierarchy generally supported the spatial dissociation implied by our *a priori* ROI analyses, where rostral motor association areas more reliably encoded transformation information (**Fig. 6a**) while caudal primary motor and visual areas more reliably encoded movement-direction information (**Fig. 6b**). Broadly, evidence of transformation-context encoding appeared to be present throughout a wider range of frontal and temporal regions (see *Discussion*).

**Figure 6.**
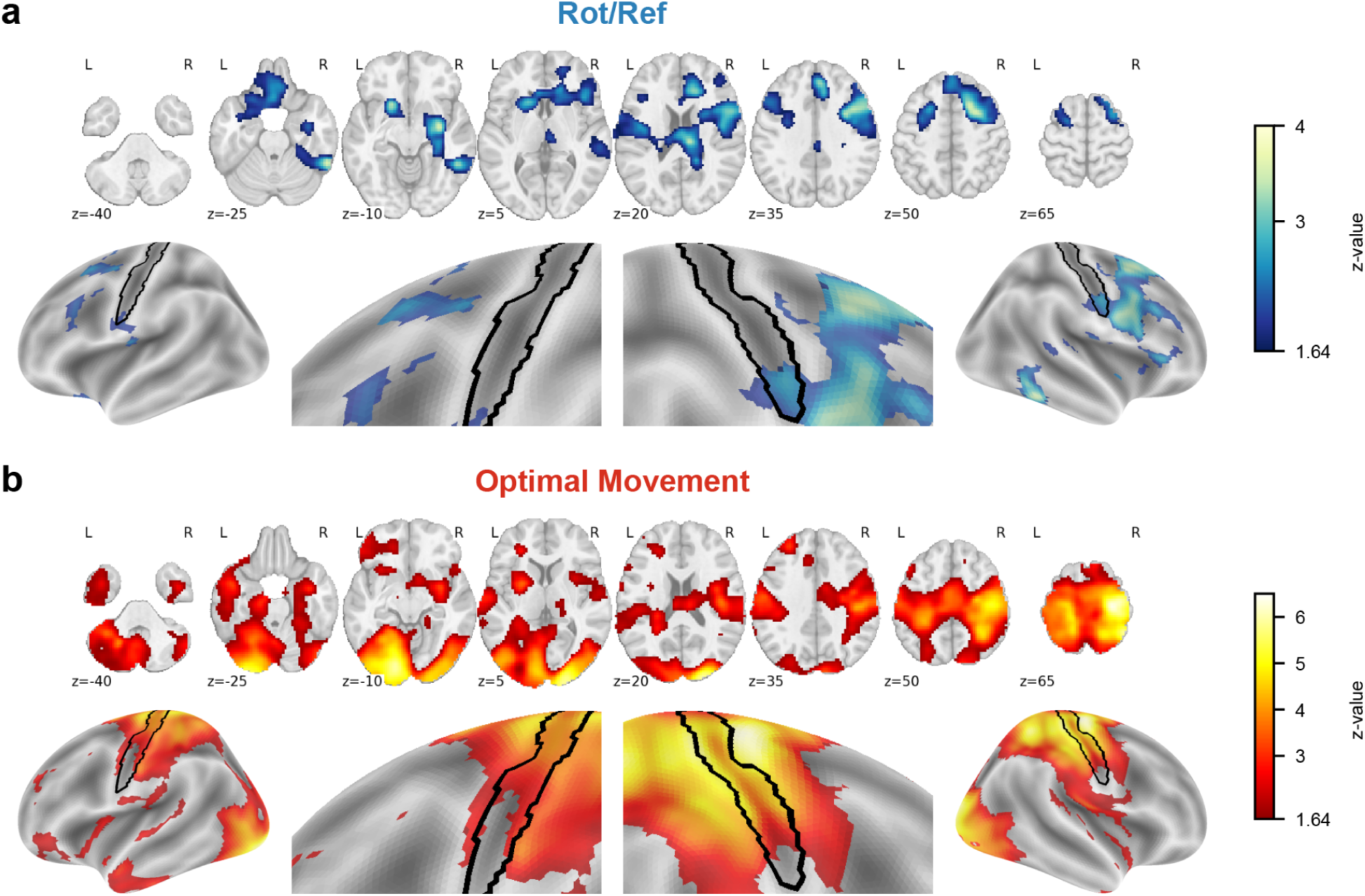
Exploratory RSA searchlight analyses. Searchlight results for **a)** transformation context encoding and **b)** movement encoding. Results are visualized on axial slices, on a zoomed-in view of sensorimotor areas surrounding the central sulcus (black outline), and on the full lateral cortical surface. Color temperature reflects voxelwise z-values. Maps are cluster-corrected at FWER *p* <.05 (with cluster-forming threshold p < .05; see *Methods* for additional details).

## Discussion

The motor system represents contextual information beyond motor commands. Where and how this information is represented will inform our understanding of how motor skills can be deployed flexibly across contexts. In this study, we used fMRI to ask if human sensorimotor cortex represents sensorimotor mappings abstracted from specific motor commands. Human participants learned to implement two different visuomotor transformations, allowing us to distinguish between neural representations of movement directions versus abstract transformation context (**Figs. 1, 2**). We found that neural activity patterns in bilateral premotor cortex encoded both kinds of information, whereas activity patterns in primary motor cortex encoded only movement direction (**Fig. 3**). These findings were robust to several potential confounds. Transformation context encoding in the left dorsal premotor cortex covaried with behavioral performance (**Fig. 4**). Moreover, the overlap between abstract context representations and lower-level movement representations was correlated with the extent to which participants employed a context-insensitive “caching” strategy (**Fig. 5**). Exploratory whole-brain analyses revealed suggestive evidence for transformation-context information across a wider set of regions (**Fig. 6**).

These findings contribute to a growing body of evidence that premotor areas encode surprisingly sophisticated abstract information for movement control. For example, invasive recordings of premotor areas in humans have revealed correlated patterns of premotor activity for functionally homologous movements, even across effectors (e.g., hand opening and closing versus toe opening and closing)^31,35^. Additionally, individuals with damage to dorsal premotor areas appear to have deficits in imitating abstract movement trajectories even when lower-level movement parameters are unaffected^28^, implicating the premotor cortex in higher-level, effector-independent action representations. Recent work in monkeys has demonstrated encoding of so-called ‘action symbols’ (recombinable representations of short action motifs that are invariant to low-level movement parameters) in ventral premotor areas^49^. These studies echo work in monkeys pointing to goal-based encoding of observed actions in premotor regions^50^. Interestingly, our results also suggest that among the brain regions encoding transformation context, the (left) dorsal premotor cortex may serve a unique role in contributing to performance during the task — a finding consistent with research that points to its role in processing the control of actions within a task, rather than general encoding of the task environment^51^.

In contrast, action representations in primary motor cortex are thought to be more concrete, a loose dichotomy that is consistent with our results here. For instance, while premotor areas deploy a more world-based coordinate frame during goal-directed movement, primary areas utilize more intrinsic, perhaps muscle-based codes^3,52^. Similarly, recent studies have shown that the primary motor cortex decorrelates contralateral and ipsilateral activity patterns^53^, in contrast to premotor areas which contain a movement coding subspace that generalizes across ipsilateral and contralateral limbs^35^. Some emerging results have complicated this view of primary motor cortex, however, showing that it can neurally segregate similar movement command representations between different task contexts^54^. Critically, this observed contextual separation of similar movement representations in primary motor cortex is robust when feedback control is required, but is less reliable in purely feedforward contexts. Our task—which involved a brief wrist-flicking movement with no time for feedback corrections—was fully feedforward, and thus our results are consistent with these emerging findings. Future work using closed-loop tasks may be useful to determine in what contexts human sensorimotor cortex ‘lumps’ or ‘splits’ representations of overlapping motor commands.

Interestingly, we found that on matched trials, where target locations and the movement solutions are matched across transformation contexts, people adopted a context-insensitive strategy (**Fig. 5**). That is, reaction times in Rotation and Reflection trials were less differentiated during matched trials compared to unmatched trials. This is consistent with prior work showing that in visuomotor rotation learning tasks, people either implement a parametric rule on each trial (i.e., gradually mentally rotating their movement plan to counteract a learned visuomotor rotation) or ‘cache’ a simpler stimulus-response strategy for oft-visited target locations (i.e., ‘reach to direction X when you see target Y’)^23,43^. These competing strategies trade off based on task demands, such that in tasks with only a few target locations, over-trained stimulus-response associations lead to the more efficient caching strategy; in contrast, in complex environments with many possible targets, people tend to compute an effortful transformation-based strategy each trial. Our study extends these effects to a new situation, where different rules have to be maintained across trials, but can be ignored on trials where both rules specify the same movement (matched trials). Our findings link these dissociable cognitive strategies to the brain, suggesting that cognitively implementing a visuomotor transformation may be driven (at least in part) by premotor areas, while producing a cached response may be driven by other brain regions.

Beyond sensorimotor cortical areas, which were our main focus, our exploratory whole-brain and additional ROI analyses revealed suggestive evidence for broader network-level effects (**Fig. 6 and S5**). First, movement-related activity patterns in our searchlight revealed robust activity in the sensorimotor network and visual cortex. But abstract transformation-context encoding painted a different picture: in addition to premotor areas, the transformation searchlight map, along with the exploratory ROI RSA analyses, highlighted a collection of regions that resembled (in broad strokes) several regions of the fronto-parietal network (FPN)^55^. This is consistent with previous fMRI work linking FPN and premotor regions to cognitive control and the maintenance of ‘task sets’ – mental representations of currently appropriate stimulus-response mappings^36,37,56–59^. Another potentially relevant finding is the implication of the premotor cortex as a node in the brain’s “physics network”, which is involved in representing predictions of physical events^60,61^. Future research could test if premotor representations related to action planning or execution, as in our task, are generalized to sensory predictions in non-motor contexts as well.

A key open question then is how movement-related task-set representations in frontal or parietal areas may interface with premotor cortex during action selection. Speculatively, it may be that rule representations in our task are even more abstract (e.g., “I’m in a mirror block”, “I’m in a rotation block”) in these areas^62^, while premotor cortex might encode the specific spatial and motor content needed to effectively select movements (e.g., “plan movement across the mirror axis”, “rotate movement plan 40° from the target”)^51^. Our caching analyses (**Fig. 5**) offer some preliminary support for this idea; while people must consistently maintain in memory the current block context on all trials (so that they can effectively react to unmatched trials), the putative reduction of premotor transformation information on matched trials suggests that it may not be simply representing a background task rule. Future experiments that vary the specific features of transformations (e.g., rotation size or sign, mirror axis location, etc.), or separate planning from execution stages, could test this idea more directly. Either way, our results are consistent with previous work in cognitive control tasks showing flexible encoding of rules in frontal areas including the premotor cortex.

Our study has several important limitations. First, we cannot definitively interpret the strong bilaterality seen in our results. Previous work has revealed ipsilateral encoding of movements in sensorimotor areas^53,63^, which could be driving our ipsilateral effects. However, even though movements of the joystick were made with the dominant (right) hand, participants simultaneously stabilized the joystick with their opposite (left) hand. Thus, ipsilateral (right hemisphere) neural results could be affected by stabilization movements (or movement plans) made with the contralateral hand. Second, our rapid event-related study design precluded us from separating planning versus execution stages of movement given the low temporal resolution of fMRI. It would be informative to see how abstract transformation encoding unfolds over time within trials, and how putatively distinct transformation versus movement neural coding axes might emerge from planning to execution stages of movement. Finally, we performed our neuroimaging on participants who had already reached a performance asymptote in this simple task; future work could target the learning process itself.

Our results reveal a potential role for premotor areas in bridging abstract sensorimotor contexts and concrete movement commands. These findings support the idea that the premotor cortex subserves the more cognitive functions involved in motor control and motor learning, such as imitation^28^, spatial and temporal abstraction of movements^49,64–68^, and perhaps the use of explicit cognitive strategies in sensorimotor learning^17,19,21^. Examining these more cognitive functions of the human motor system will help us better understand the computational and neural coding principles that give rise to the impressive range and flexibility of human action.

## Methods

### Participants

Thirty-one right-handed people (aged between 18-35, no history of neurological disease or injury, fluent in English and normal or corrected-to-normal visual and auditory acuity) participated in the study. Handedness was assessed using the Edinburgh Handedness Inventory (EHI)^69^. All procedures were approved by the Yale Institutional Review Board, and participants were paid $30 US dollars per hour of participation.

Two participants did not complete their sessions because the tutorial took too long (N = 1) or because of technical issues with the scanner (N = 1). Additionally, two participants’ data were excluded from analysis due to head motion exceeding our predefined threshold (fewer than six runs left after excluding runs with more than 70 rejected volumes after applying “optimized scrubbing”^70^, with framewise displacement (FD) threshold of 0.5mm, and standardized DVARS threshold of 1.5). As a result, we used data from 27 participants for analyses (18 F; age, 18-32; mean, 23.7; SD, 3.9 years; EHI score M = 89.0, SD = 19.0).

### Task Procedure

Participants used a joystick during the training phase outside of the scanner (Logitech Extreme 3D Pro) and while inside the scanner (Tethyx) to perform ballistic wrist movements to guide a cursor onto a circular target goal (radius of 6mm). The target location on a given trial was chosen from four possible positions, placed 20° from the central vertical line clockwise or counter-clockwise in either the upper or lower half of the workspace, at a distance of 10cm from the center position. Target location was pseudo-randomized across trials such that no target position was shown twice in a row within a block. Participants controlled a small visual cursor with the joystick (red circle, 3.1mm radius). On each trial, participants started from the center position of the screen (circle with 5mm radius), which corresponded to the neutral resting position of the joystick.

Each trial started after a variable inter-trial interval (ITI) of 1-4s. Then participants were asked to maintain the joystick at a neutral resting position, and only the center circle appeared on the screen (“Return” phase). Afterward, the target appeared on the screen with the center hollowed out. Participants were told not to move during this period, but to plan their upcoming movement (“Wait” phase, length sampled from a uniform distribution from 2 s to 4 s). If participants moved their cursor beyond the circular start area (2.5cm radius around the center position of the screen), the trial aborted and a “HOLD STILL” message was displayed for 3s, after which they repeated the same trial again. Following the Wait phase, the target was filled in, cueing participants to move their joystick as quickly as possible to guide the cursor to the target, with a movement time constraint of 1s (“Move” phase). If participants did not complete their movement within the 1s limit, they would see a “TOO SLOW!” error message for 3s, after which the task would move on to the next trial. The Move phase was followed by a 1s delay before the terminal cursor position was displayed (for 1s) as feedback. Feedback presentation was delayed to limit implicit motor adaptation and isolate explicit visuomotor learning^23,38^.

Crucially, terminal cursor positions could be either rotated clockwise by 40° relative to the movement angle, or reflected about a vertical mirror axis. Participants were told what kind of transformation (Rotation or Reflection) they would experience at the start of each block, and experienced that same transformation throughout the block. Each block consisted of 16 trials, with no consecutive repeating trials with the same target location. To train participants on the task, we had them perform a tutorial version of the task outside of the scanner. In this tutorial phase, participants first practiced joystick reaches without any transformations for one block. Afterwards, they experienced the rotation and the reflection transformation for one block each, in that order. The transformations were explicitly explained to participants via verbal instruction and a demonstrational video, helping them to perform the task effectively prior to scanning. The tutorial learning blocks were repeated until participants reported that they sufficiently understood the instructions.

After the tutorial, participants performed the task again inside the MRI scanner. During their anatomical scan, they first performed one block of practice with no transformation to familiarize them with the change in setup (i.e., laying supine, and using the MR-compatible joystick). Then, they practiced the standard task again under either rotation or reflection for one block each. During the functional runs, they performed one block of each transformation (with order varying across runs, such that participants experienced five rotation-first runs and four reflection-first runs total, but in shuffled order) per run for nine runs (eight for one participant). The main task during functional scanning thus consisted of 288 total trials (**Fig. 1c**). After task completion, participants were taken out of the scanner and compensated for their time.

### Behavioral Analysis

All behavioral analyses were performed using R (version 4.2.0)^71^. To quantify task performance, we analyzed the joystick coordinate data (sampling rate: 60Hz) to obtain two measures of error for each trial: initial error and endpoint error. Initial error was quantified as the angle of the segment of the movement between 1cm-3cm from the center position, and endpoint error was computed as the angle of movement at the full radial distance of the workspace (i.e., from the center position to the invisible ring containing the targets, 10cm). For error estimation, all trajectories were rotated such that the target location was normalized to 0°. To estimate movement speed, we applied smoothing using a second-order Savitzky-Golay^72^ filter (filter length: 7 frames) on the trajectory data using the *signal* package^73^. Then, we calculated the time following the go-cue at which the speed of framewise movement exceeded 2cm/s. This point was taken as the reaction time (RT), and the remaining time it took for the movement to exceed the radial distance of the workspace was taken as the movement time (MT). For visualization of reaching movements in **Fig. 2d**, we used the smoothed trajectories in each of the eight trial types from an example participant to visualize trajectories reflecting the correct hand reach angle.

We tested whether there were significant differences in performance between the Rotation and Reflection conditions both for initial and endpoint angular error, as well as RT and MT measures, using paired t-tests. To quantify whether or not learning persisted into the fMRI phase of the task, we estimated the linear slope of error over scanning trials per each condition and subject, and then tested whether the slopes per condition were significantly different from zero via one-sample t-tests.

Additionally, we asked if people learned the two transformations comparably between the two conditions during the tutorial learning phase (which also included explicit instructions about the perturbations). To do this, we compared the initial heading error in both Rotation and Reflection trials in the first two versus the last two trials of the learning. Importantly, for visualization and statistical purposes, we flipped the sign of the error for the two Reflection trial types (the two unmatched targets, where the optimal hand angle was in the opposite direction of the other trial types) such that higher error would always mean the participant is making an error in the same direction away from the optimal movement. To assess whether participants arrived at asymptote performance in later learning, we derived linear slopes for participants’ errors in both conditions in the latter half of the learning (trials 9-16), and then compared it to zero using a one-sample t-test. In case the learning phase was performed more than once, we visualized the data from their latest attempt at the learning phase.

### Neuroimaging Data Acquisition

A 3T Siemens Magnetom scanner with a 64-channel head coil was used to perform anatomical and functional scans. For the anatomical scan, a T1-weighted MPRAGE (magnetization-prepared rapid acquisition gradient echo) sequence was used (TR: 2400 ms; TE: 2.19 ms; field of view (FOV): 256 x 256 mm; flip angle: 8°; orientation: sagittal; slices: 208; voxel sizes: 1 x 1 x 1 mm). Functional images were collected with a gradient echo planar imaging sequence (TR: 1058ms; TE: 30ms; field of view (FOV): 225 x 225 mm; flip angle: 57°; orientation: axial; slices: 60; voxel size: 2.5 x 2.5 x 2.5 mm; multiband acceleration factor: 4). Two fieldmaps were acquired, one before the first functional imaging sequence and another before the fifth. They were acquired with identical imaging parameters (TR: 7033 ms, TE: 80 ms, orientation: axial, flip angle: 90°; slices: 60), to account for magnetic field inhomogeneities.

### Preprocessing

Preprocessing of the fMRI data was performed using fMRIPrep 23.2.1^74^, which is based on Nipype 1.8.6^75^, as follows.

#### Anatomical data preprocessing

The T1w image was corrected for intensity non-uniformity (INU) with N4BiasFieldCorrection (Tustison et al.^76^), distributed with ANTs 2.5.0 (Avants et al.^77^, RRID:SCR_004757), and used as T1w-reference throughout the workflow. The T1w-reference was then skull-stripped with a *Nipype* implementation of the antsBrainExtraction.sh workflow (from ANTs), using OASIS30ANTs as target template, with the exception of one participant, whose brain was extracted using FSL’s Brain Extraction Tool (BET) with fractional intensity threshold of .5. Brain tissue segmentation of cerebrospinal fluid (CSF), white-matter (WM) and gray-matter (GM) was performed on the brain-extracted T1w using fast (FSL, RRID:SCR_002823, Zhang, Brady, and Smith^78^). Volume-based spatial normalization to MNI152NLin2009cAsym space was performed through nonlinear registration with antsRegistration (ANTs 2.5.0), using brain-extracted versions of both T1w reference and the T1w template. The following templates were selected for spatial normalization and accessed with *TemplateFlow* (23.1.0)^79^: *ICBM 152 Nonlinear Asymmetrical template version 2009c*^80^.

#### Functional data preprocessing

For each of the BOLD runs found per subject (across all tasks and sessions), the following preprocessing was performed. First, a reference volume was generated, using a custom methodology of *fMRIPrep*, for use in head motion correction. Head-motion parameters with respect to the BOLD reference (transformation matrices, and six corresponding rotation and translation parameters) are estimated before any spatiotemporal filtering using mcflirt (FSL , Jenkinson et al.^81^).

The estimated *fieldmap* was then aligned with rigid-registration to the target EPI (echo-planar imaging) reference run. The field coefficients were mapped on to the reference EPI using the transform. The BOLD reference was then co-registered to the T1w reference using *bbregister* (FreeSurfer) which implements boundary-based registration (Greve and Fischl82). Co-registration was configured with six degrees of freedom.

Several confounding time-series were calculated based on the *preprocessed BOLD*: framewise displacement (FD), DVARS and three region-wise global signals. FD was computed using two formulations following Power (absolute sum of relative motions, Power et al.^70^) and Jenkinson (relative root mean square displacement between affines, Jenkinson et al.^81^). FD and DVARS are calculated for each functional run, both using their implementations in *Nipype* (following the definitions by Power et al.^70^). The three global signals are extracted within the CSF, the WM, and the whole-brain masks.

Additionally, a set of physiological regressors were extracted to allow for component-based noise correction (*CompCor*^82^). Principal components are estimated after high-pass filtering the *preprocessed BOLD* time-series (using a discrete cosine filter with 128s cut-off) for the two *CompCor* variants: temporal (tCompCor) and anatomical (aCompCor). tCompCor components are then calculated from the top 2% variable voxels within the brain mask. For aCompCor, three probabilistic masks (CSF, WM and combined CSF+WM) are generated in anatomical space. The implementation differs from that of Behzadi et al. in that instead of eroding the masks by 2 pixels on BOLD space, a mask of pixels that likely contain a volume fraction of GM is subtracted from the aCompCor masks. This mask is obtained by thresholding the corresponding partial volume map at 0.05, and it ensures components are not extracted from voxels containing a minimal fraction of GM. Finally, these masks are resampled into BOLD space and binarized by thresholding at 0.99 (as in the original implementation). Components are also calculated separately within the WM and CSF masks. For each CompCor decomposition, the *k* components with the largest singular values are retained, such that the retained components’ time series are sufficient to explain 50 percent of variance across the nuisance mask (CSF, WM, combined, or temporal). The remaining components are dropped from consideration. The head-motion estimates calculated in the correction step were also placed within the corresponding confounds file. The confound time series derived from head motion estimates and global signals were expanded with the inclusion of temporal derivatives and quadratic terms for each^83^. Frames that exceeded a threshold of 0.5 mm FD or 1.5 standardized DVARS were annotated as motion outliers.

### First-level General Linear Model

All neuroimaging analyses were performed using Python (version 3.10.13 for all analyses except for multidimensional scaling (MDS), which used 3.11.5). Preprocessed run-wise BOLD images were used to fit a first-level GLM in each scanning run, in order to derive a neural activation map for each trial.

To do this, trial-wise regressors were first constructed, modeling BOLD activity over the interval between each trial’s “Wait” onset (**Fig. 1b**) through to the end of the trial. Additionally, the first six aCompCor components as well as the six head motion regressors derived during preprocessing were extracted and added as nuisance regressors. To account for low-frequency signal drift during the run, five cosine drift regressors derived from fmriPrep were also included as nuisance regressors. We used the resulting design matrix along with an ar1 noise model and the canonical double-gamma hemodynamic response function to fit our first-level GLM using Nilearn^84^’s *FirstLevelModel* function. Trial-wise activation maps were estimated as beta coefficients for each trial regressor, and each voxel’s trial-wise activation estimates were z-scored across trials within each run. The resulting activation maps were used for all multivariate analyses. Aborted trials were not included in the model. Voxel-wise time series were standardized within run before the GLM was fit.

### Univariate Contrast

We performed a whole brain group-level contrast between conditions (Rotation vs. Reflection) using the trial-level activation maps obtained from the first-level GLM. To do this, we first averaged trial-wise whole-brain activation maps for each condition for each participant. Then, for each run, we calculated the difference between the averaged activation map for the Rotation and the Reflection conditions. Finally, this difference map was averaged across runs to produce one contrast map per participant. After smoothing at 6mm FWHM Gaussian kernel, the images were used to perform a two-sided one-sample t-test. This was done using Nilearn’s *non_parametric_inference* function, to implement FWER permutation-based cluster correction (cluster forming threshold p < .05; cluster mass threshold p < .05) over 10,000 permutations.

### Representational Similarity Analysis

In our primary neural analysis, we investigated which task features were represented in activity patterns in our *a priori* ROIs. We first obtained anatomical masks for these regions using the Julich brain atlas^85^. We obtained maximum probability maps (MPM) for the following regions: dorsal and ventral premotor cortex^86^, primary motor cortex^87^, dorsolateral prefrontal cortex^88^, lateral occipital cortex^89^, primary somatosensory cortex^90,91^, hippocampus^92,93^, and the superior parietal lobe^94,95^.

Next, for each participant, we obtained a cross-validated empirical correlation matrix to estimate, in an unbiased manner, activity patterns across trial types within each region. To do this, we averaged the region activation maps for the eight unique trial types (4 target locations X 2 conditions) within each run. Then, we performed Spearman correlations between each of the eight trial types. The correlations were computed using split-half cross-validation over odd versus even runs. Thus, each element in the correlation matrix represented the correlation between the average activation map in odd versus even runs. This resulted in a 8x8 unbiased empirical correlation matrix per participant and ROI that was symmetric across the diagonal.

Afterwards, we modeled this matrix as a linear combination of hypothesized task-based correlation matrices, as depicted in **Fig. 3a**, including intercept. The model was fit by ordinary least squares using the 36 elements on the upper triangle of each matrix, including the diagonal. Each of the model matrices exemplified a candidate representation that might be present in a given brain region, assuming perfect correlation between trial types that shared each task feature. As a result, we obtained participant- and ROI-level regression coefficients for each of the model matrices. We performed a one-sample, one-tailed t-test against zero for each ROI for each model to examine which regions exhibited significant positive evidence of activity patterns containing information related to each candidate representation. Given the large number of *a priori* ROIs, the results of all t-tests on these beta weights were FDR-corrected. Additionally, we tested for relative encoding of transformation-context information versus optimal movements by performing a repeated-measures ANOVA on the regression coefficient differences, with factors brain region (premotor versus primary motor, with separate models for dorsal and ventral premotor) and hemisphere.

We performed a participant- and ROI-wise model comparison between the regression models that either included the transformation context factor (‘full’ model), or that did not (‘reduced’ model), with the intercept included. We then compared the two models’ adjusted R-squared and calculated the change in AIC (Akaike Information Criterion) values summed across participants, as visualized in **Fig. S1**. For comparison, we calculated the ‘noise ceiling’ adjusted R-squared to estimate the amount of variance there is within the empirical correlation matrix being modeled that can be explained by any regression model. Specifically, it was estimated by modeling a single participant and ROI’s empirical correlation matrix with the average empirical correlation matrix of all participants including the participant being modeled, in the same ROI^96^. This ceiling is represented as the horizontal line in **Fig. S1a**. Delta AIC was calculated by subtracting the summed AIC of the full model from the reduced model, such that positive values would indicate support for the full model.

We also performed an exploratory searchlight analysis to visualize results across the whole brain. Specifically, for each voxel within the brain mask, we constructed a small spherical searchlight with a radius of 5 voxels, using the rsatoolbox^97^ *get_volume_searchlight* function. Then, we iterated over each searchlight fitting the same regression model as above. We attributed the resulting regression coefficients to the voxel at the center of each searchlight. We included searchlights that at least included half of the voxels within the brain mask in the searchlight analysis. We note that the size and shape of the searchlights was not optimized to match our ROI results or any *a priori* predictions or anatomical features. We visualized the resulting group t-maps on axial slices of the MNI152NLin2009cAsym template. We also projected the t-maps on the fsaverage5 cortical surface by averaging samples between white and pial surfaces (**Fig. 6**). Additionally, we used the central sulcus label in the Destrieux atlas^98^ to visualize the central sulcus for anatomical reference. The subject-level searchlight maps were smoothed with a 6mm FWHM Gaussian kernel before the group-level test. These maps were then tested with a one-sided sign-flip permutation test (10,000 permutations) with cluster-mass FWER correction (cluster forming threshold: p <.05; FWER p < .05). For display, t-values were converted to z-values, and voxels in surviving clusters were shown in **Fig. 6**. The cerebellum portion of this searchlight was masked and plotted on a cerebellar flat map^99^ (**Fig. S6**). The cerebellum mask used in this analysis was first obtained in MNI152NLin6AsymC space as an atlas from the DiedrichsenLab Github repository^100^, and then transformed into MNI152NLin2009cAsym using the composite transform obtained from TemplateFlow^79^. The transformation was performed using ANTspy^101^’s *antsApplyTransforms* function. The resulting cerebellar searchlight map was unthresholded for visualization.

To visualize the representational geometry of particular ROIs, we performed multidimensional scaling (MDS) on empirical correlation matrices. We first averaged empirical correlation matrices to obtain a group-level correlation matrix. After calculating a dissimilarity matrix by subtracting this matrix from 1, we passed the resulting dissimilarity matrix to the scikit-learn^102^ implementation of classical MDS, with diagonals converted to zero.

We also performed an exploratory brain-behavior correlation analysis, testing for an association between the neural representation of transformation context (Rotation vs. Reflection) and behavior. To do this, we correlated regression coefficients for the Rotation/Reflection predictor from our RSA (see above) with behavior, using the average absolute angular error per participant as an index of overall performance. We tested for this correlation in our premotor ROIs as they were associated with reliable positive coefficients for Rotation/Reflection encoding in the RSA; primary motor cortex was used as a control ROI for comparison.

Finally, we performed an additional control analysis asking if neural encoding of transformation context was confounded with motor behavior in a manner not accounted for by our primary RSA regression. To do this, first we modeled the same RSA regression as before, but substituted the optimal movement matrix with an empirical angular similarity matrix, where each element represents the mean initial hand angle similarity between trial types. We next utilized initial cursor reach angle, which should not correlate with transformation context by design, to further confirm if context information is discoverable in behavioral data in a way that might be confounded by what we observe in neural analyses. We constructed a cross-validated (split-half) “behavioral similarity matrix” across the trial types. Similarity was measured as cosine similarity between mean initial cursor directions in different trial types. We then performed the same RSA regression on these behavioral data to get a behavior-derived Rotation/Reflection transformation beta value, and asked if this effect was related to the neural Rotation/Reflection transformation effect by performing linear regression. We standardized the empirical behavioral and neural similarity matrices, as well as the model matrices, such that magnitude of the slope would be interpretable. Matrices were not standardized across subjects, so the intercept is still interpretable as the presence of neural information when the behavioral information is accounted for. The logic here was that if one of our key neural RSA regression results – the neural Rotation/Reflection transformation effect – was actually driven by unmodeled behavioral distinctions between the transformation contexts, then these brain-behavior relationships (measured as slopes in the linear regression models) would be reliably positive. On the other hand, if the slopes were null and the intercepts of this regression remained significantly positive, it would suggest that behavioral effects were not a driving confound. We tested for these relationships across premotor and primary motor ROIs.

### Neural Overlap Analysis

To quantify the extent to which neural representation of transformation context and optimal movement overlap in neural patterns of interest, we estimated the angular distance between the vectors that contained information regarding these task features. Specifically, we calculated the contrast between the average neural activation patterns for all trial types in the Rotation condition versus the Reflection condition, from which we derived the ‘Rot/Ref’ vector. This way, target location was balanced across both sides of the contrast, while optimal movement added up to zero, under an idealized assumption of approximate cosine-like tuning of brain activity patterns to movement direction^103,104^. Next, to derive the ‘Movement’ vector, we used a contrast between the lower right target location in the Rotation condition (Rotation 4) plus the upper left target location in the Reflection condition (Reflection 2), and the average between upper left target location in the Rotation condition (Rotation 2) and the lower right target location in the Reflection condition (Reflection 4). This way, the target location information and transformation context information was balanced across both sides of the contrast, while there was a remaining imbalance in the average optimal movement direction. The logic of these analyses yields a vector in which we expect transformation information to be encoded, and another in which we expect optimal movement direction information to be encoded.

Next, we obtained the cosine angle between the Rot/Ref and Movement vectors described above. To avoid bias in estimation due to noise we performed a split-half correlation, where we obtained both vectors from the neural activation patterns across trials in odd and in even runs separately. Then, we obtained cosine angles between the Rot/Ref vector within odd runs and Movement vector in even runs, and the Rot/Ref vector within even runs and Movement vector in odd runs. We took the product of these two cosine angles as the neural overlap measure, such that positive estimates would indicate larger overlap (i.e., more acute or more obtuse angles). That is, if the angle between was either smaller than or larger than 90 degrees between the two vectors, our overlap measure would indicate some degree of positive overlap between the two components (since the sign of direction is arbitrary in our contrast; the Rot/Ref vector would contain transformation context information if we defined the vector as Rotation minus Reflection or Reflection minus Rotation). Finally, we estimated the relationship between this neural overlap measure, in either the left premotor cortex or the left primary motor cortex ROIs, with a behavioral ‘caching index’ for each participant (see *Results*). The caching index was derived by subtracting the absolute RT difference between conditions in matched trials from unmatched trials. Here, positive values would indicate more behavioral ‘caching’ of performance between Rotation and Reflection conditions in matched trials (i.e., more ‘similar’ behavioral signatures), where optimal movement and target location are matched across transformations.

## Acknowledgments

The authors are supported by the National Institute of Neurological Disorders and Stroke (1R01NS132926-01 to S.D.M.). This study was also supported by BrainWorks at the Center for Neurocognition and Behavior in the Wu Tsai Institute, Yale University *(RRID:SCR_024556)*.

## Declaration of Interests

The authors declare no competing interests.

## Notes

### Competing Interest Statement

The authors have declared no competing interest.

### Summary of Updates

searchlight figures and univariate contrast figure updated; methods and manuscript updated to be consistent with changes.

