## Supplemental Materials for "Abstract Representations of Sensorimotor Transformations in Human Premotor Cortex"

### Supplemental Information

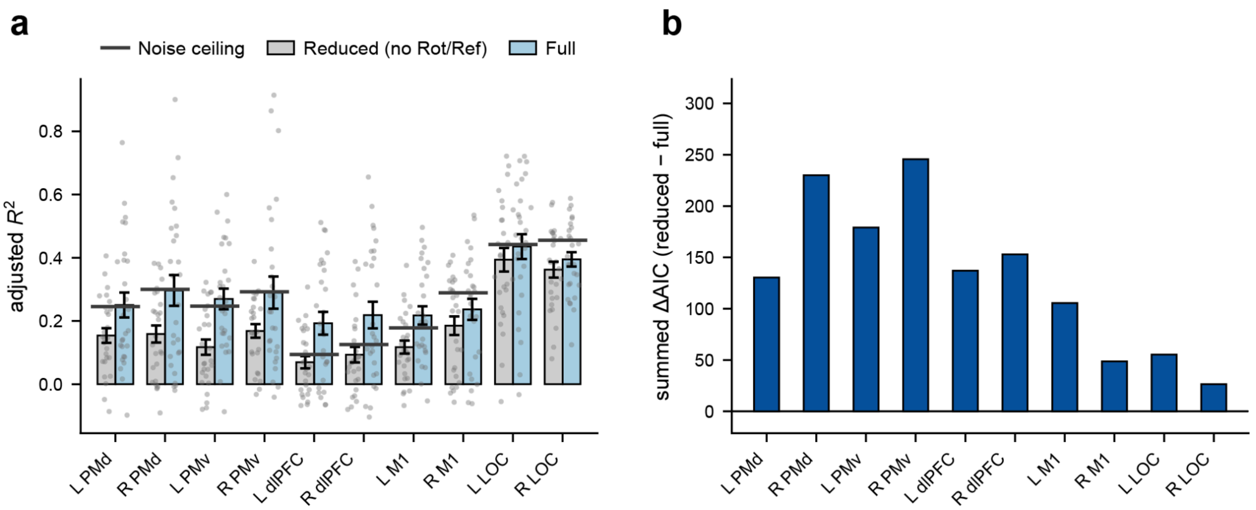


**Figure S1.** *Model comparison results between the full and reduced regression models, using adjusted R-squared (**a**) or AIC (**b**) metrics.* Error bars = 1 s.e.m.

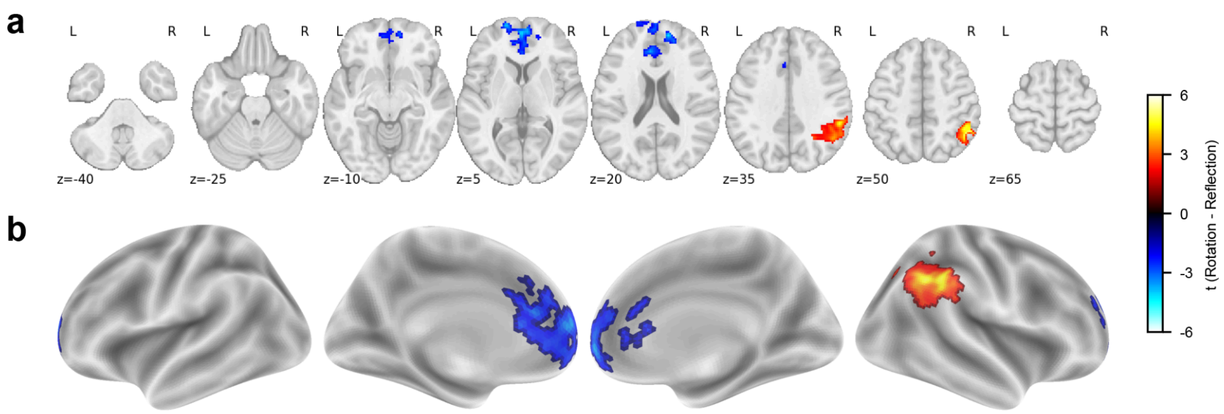


**Figure S2.** *Univariate contrast results across the whole brain.* Positive values indicate larger activation during Rotation trials relative to Reflection trials, while negative values indicate the opposite. Results are cluster-corrected at FWER $p$ <.05 (with cluster-forming threshold $p < .05$), and visualized in both **(a)** axial slices in volume space, and **(b)** the cortical surface.

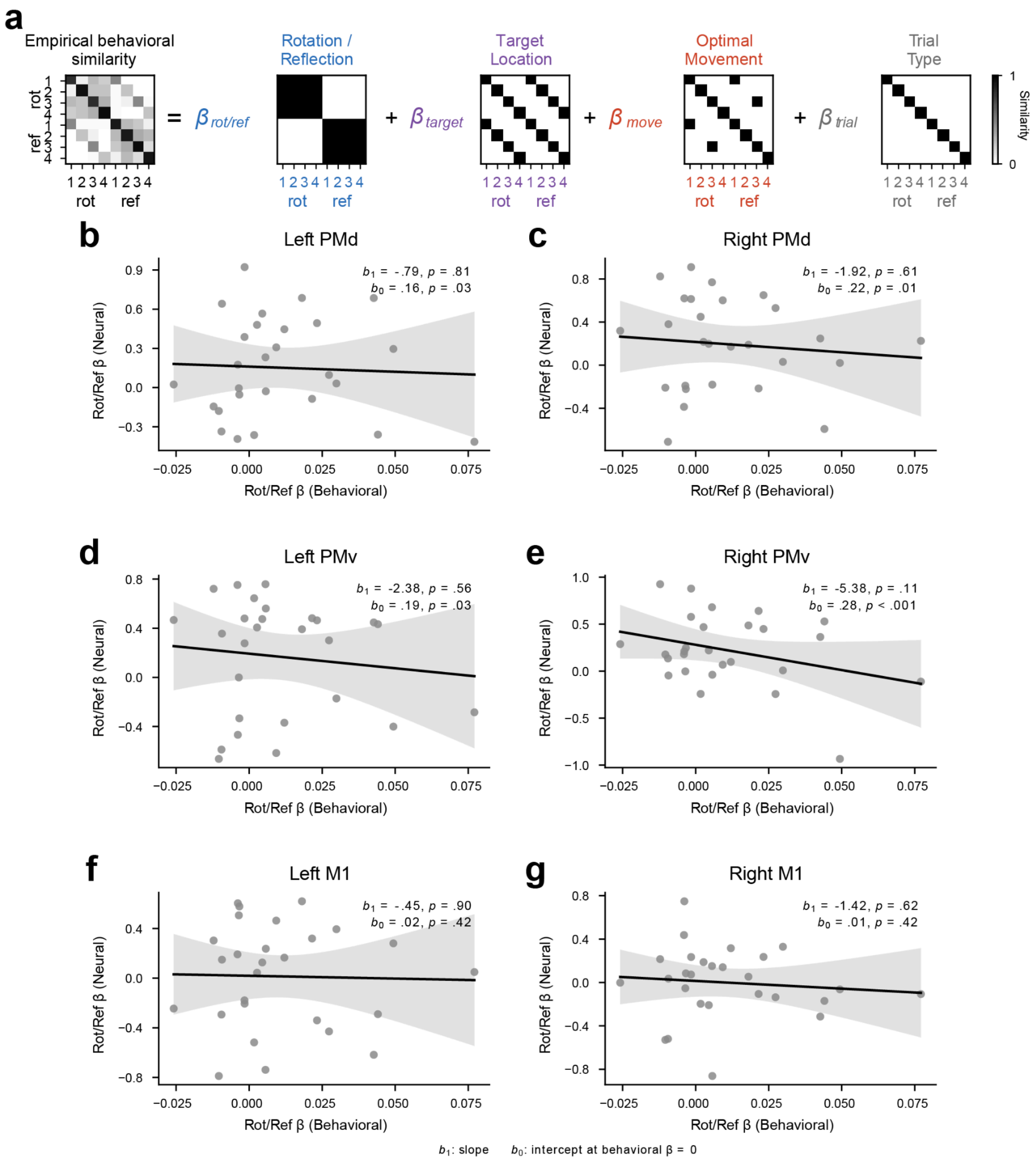


**Figure S3.** *Comparison of neural and behavior RSA regression coefficients.* **a)** In this control analysis, we computed a cross-validated empirical *behavioral* representational similarity matrix (RSM) on participants' initial cursor heading directions across conditions (cross-validated in the same manner as the neural RSM; see *Methods*). We then performed the same RSA on the behavioral RSMs as we performed on the neural RSMs (Fig. 3, main text). Finally, we extracted the beta values for the transformation-context effect and compared those across behavioral and neural data sets using linear regression. **b-g)** The behavioral-neural comparisons were done across premotor and primary motor ROIs. Insets show two-tailed one-sample t-test results on the slopes ($b_1$) and one-tailed one-sample t-test results for intercepts ($b_0$) resulting from the regressions. We note that non-significant slopes and significant positive intercepts, as seen in the premotor areas, are consistent with abstract transformation context effects that are unconfounded with movement differences across conditions. PMd = dorsal premotor cortex; PMv = ventral premotor cortex; M1 = primary motor cortex.

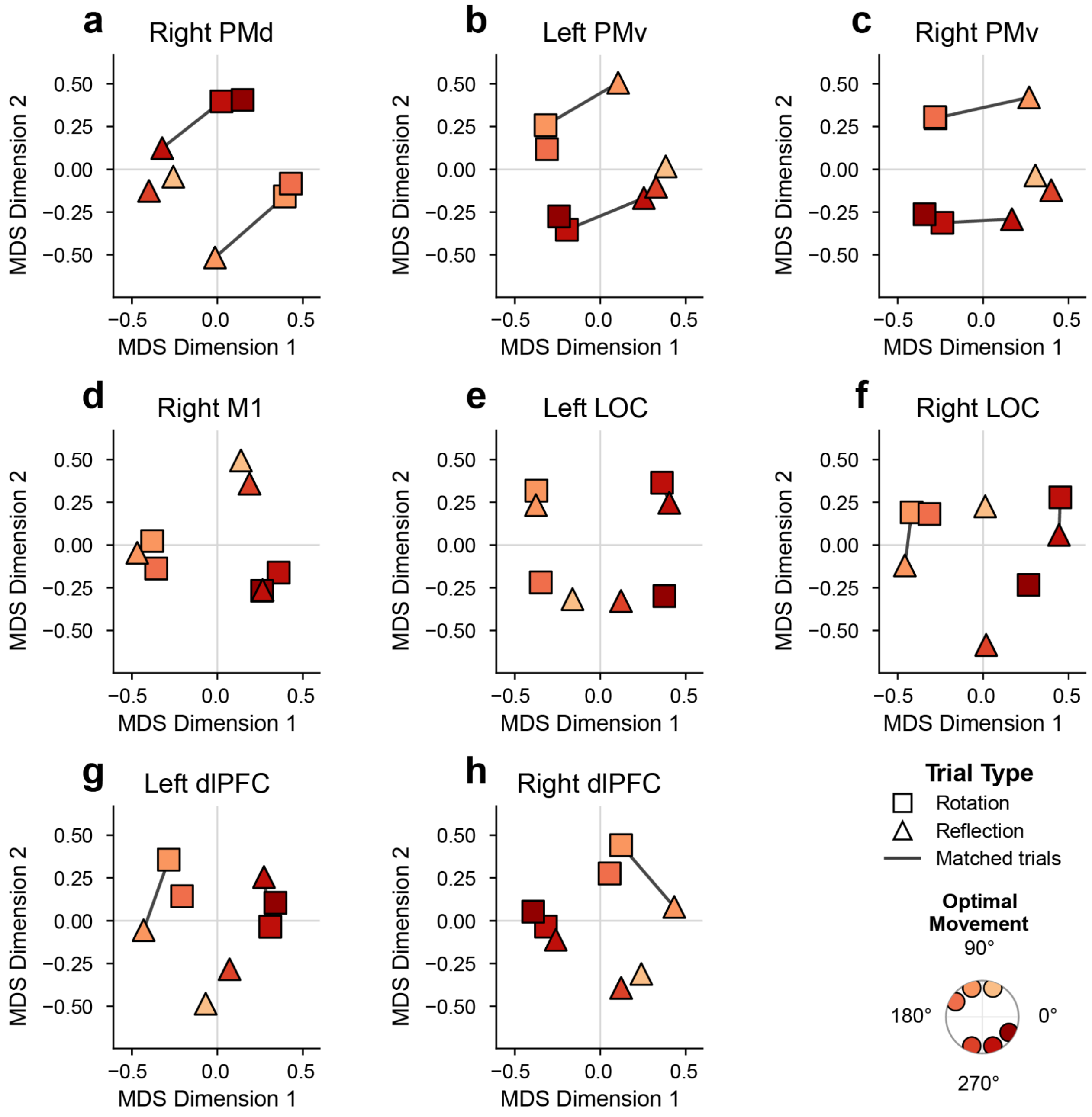


**Figure S4.** *MDS plots for additional ROIs*. **a-h**) MDS plots as in Fig. 4a, b in the main text, for our additional *a priori* anatomical ROIs (see *Methods* for further analysis details). PMd = dorsal premotor cortex; PMv = ventral premotor cortex; M1 = primary motor cortex; LOC = lateral occipital cortex; DLPFC = dorso-lateral prefrontal cortex.

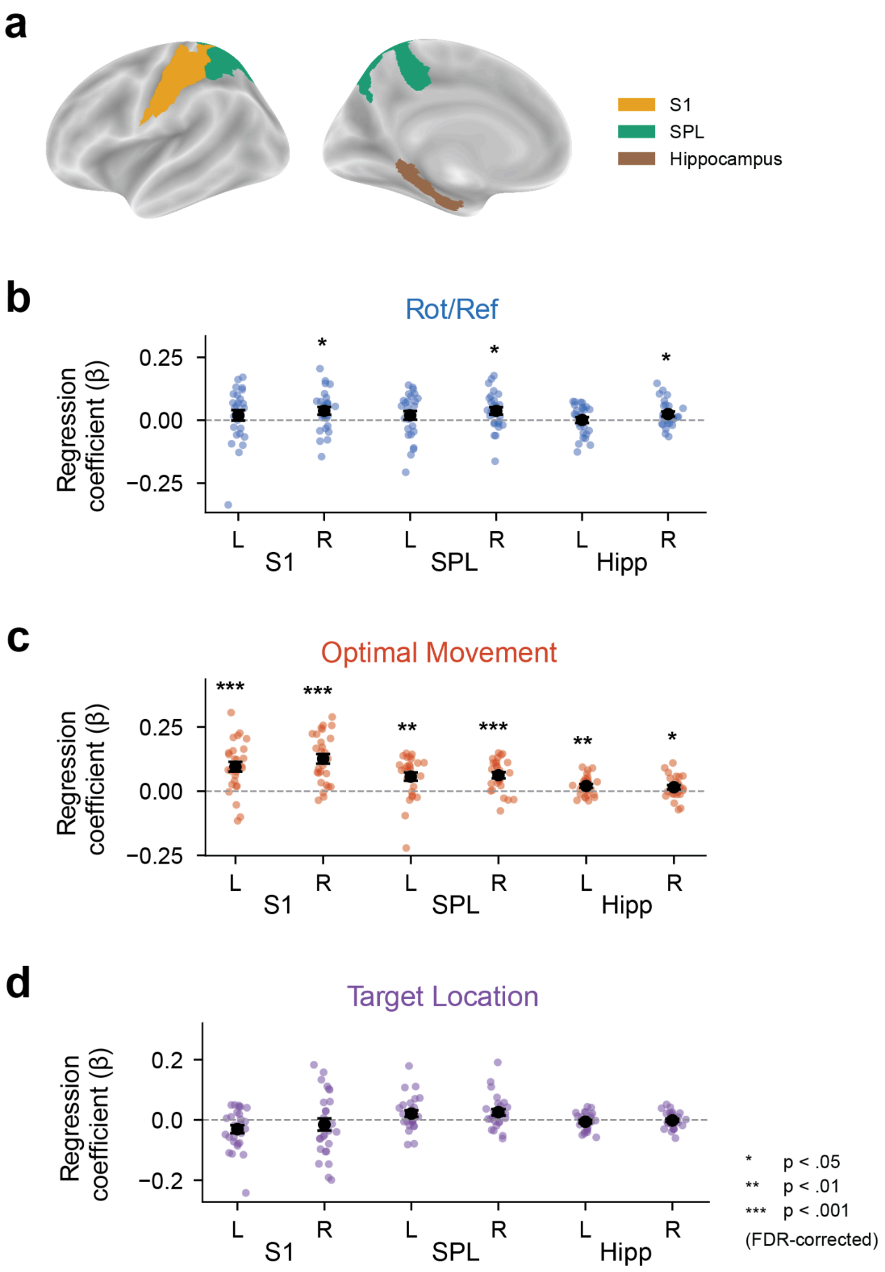


**Figure S5.** *RSA regression coefficients for additional ROIs.* **a)** Cortical surface display of additional anatomical ROIs tested in our RSA analysis, which included bilateral primary somatosensory cortex (S1), superior parietal lobule (SPL), and the hippocampus (Hipp). RSA regression coefficients for the additional ROIs, related to **b)** transformation context, **c)** movement direction, and **d)** visual target location. All t-tests on regression coefficients are FDR-corrected. Error bars represent 1 S.E.M.

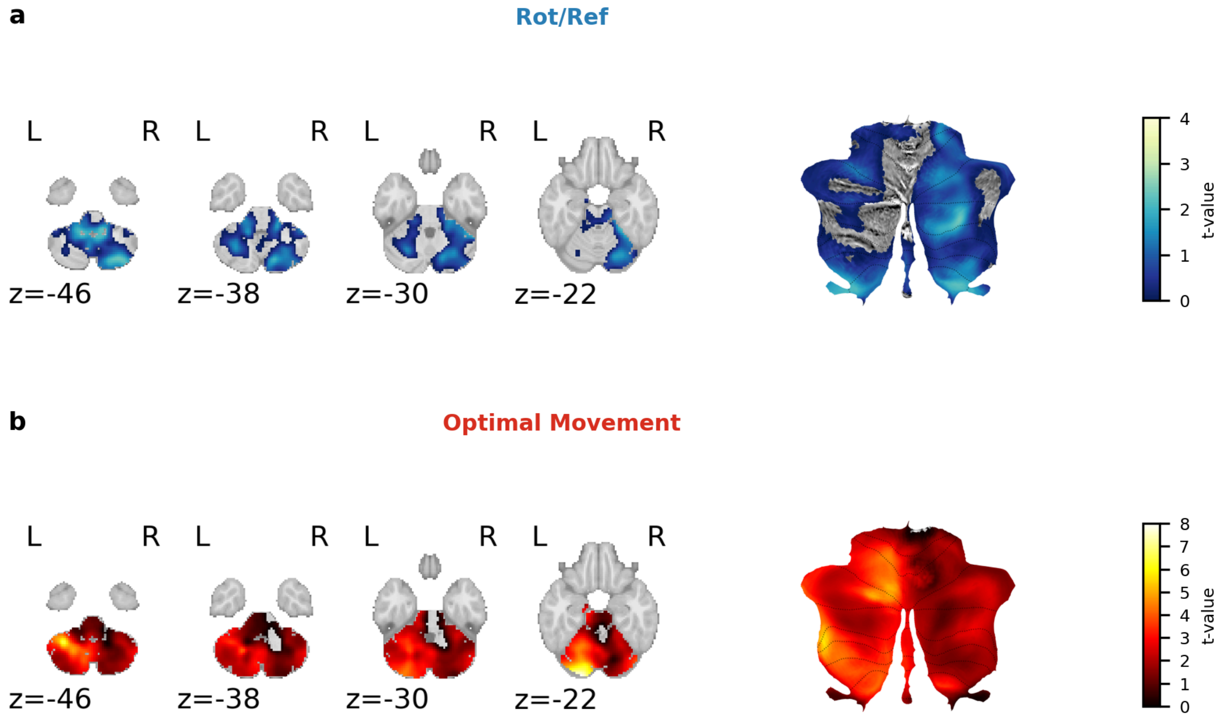


**Figure S6.** *RSA searchlight in the cerebellum.* In addition to the cortical searchlight results (Fig. 6, main text) we also visualized searchlight results in the cerebellum, both in the volume (left) and on a cerebellar 'flatmap'[99] (right), both for **a)** transformation context encoding, and **b)** optimal movement encoding. The map is unthresholded (no clusters survived correction), with the colorbars on the right representing t-values from the one-sample t-test.
